# Non-cell autonomous OTX2 transcription factor regulates anxiety-related behavior in the mouse

**DOI:** 10.1101/710848

**Authors:** C. Vincent, J. Gilabert-Juan, R. Gibel-Russo, D. Alvarez-Fischer, M.-O. Krebs, G. Le Pen, A. Prochiantz, A.A. Di Nardo

## Abstract

The Otx2 homeoprotein transcription factor is expressed in the dopaminergic neurons of the ventral tegmental area, a mesencephalic nucleus involved in the control of complex behaviors through its projections to limbic structures, including the ventral hippocampus, amygdala, nucleus accumbens and medial prefrontal cortex. We find adult mice heterozygous for *Otx2* show anxiolysis-like phenotype in light-dark box and elevated plus maze paradigms. However, the number of dopaminergic neurons, the integrity of their axons, their projection patterns in target structures, and the amounts of dopamine and dopamine metabolites in targets structures were not modified. Because OTX2 is expressed by the choroid plexus, secreted into cerebrospinal fluid and transferred to parvalbumin interneurons of the cortex, hippocampus, and amygdala, we investigated if this phenotype could result from the decreased synthesis of OTX2 in the choroid plexus. Indeed, the anxiolysis-like phenotype was partially recapitulated in the *Otx2*^+/*AA*^ and *scFvOtx2^tg/0^* choroid-plexus dependent non-cell-autonomous OTX2 loss of function mouse models. Furthermore, the phenotype was reversed by the overexpression of *Otx2* specifically in choroid plexus of adult *Otx2* heterozygous mice. Taken together, OTX2 synthesis by the choroid plexus followed by its secretion into the cerebrospinal fluid is an important regulator of the anxiety phenotype in the mouse.

## INTRODUCTION

Homeoprotein (HP) transcription factors are expressed during development inside and outside the nervous system (1). In the developing nervous system, HPs are more specifically involved in the segmentation of the neuroepithelium, cell lineage decisions, cell shape and migration, and axon guidance (2). Interestingly, many HPs remain expressed in the adult nervous system, although often in regions and cell types that differ from those identified during development (3). As such, specific HP adult physiological functions may differ and indeed remain largely uncharacterized. Nonetheless, some HPs have been shown to transfer between cells in both embryonic and postnatal contexts. This intercellular transport results in non-cell autonomous HP regulation of gene expression and possibly of the epigenetic status of the receiving cells (4). The sequences responsible for HP internalization and secretion are highly conserved (5,6), and 150 of 160 recently tested HPs were found to transfer both *in vitro* and *in vivo* (7), strongly suggesting that transduction is a shared property of most of the ~300 members of the HP family.

The vertebrate *orthodenticle* orthologue *Otx2* HP acts as a gap gene in early development for the anterior part of the central nervous system, including the telencephalon (8,9). However, *Otx2* expression is rapidly turned off and has already ceased in the mouse cerebral cortex parenchyma at embryonic day 15 (E15). In contrast, its expression persists throughout life in the pineal gland, choroid plexus (ChP), and in neurons within the septum, thalamus, cerebellum and midbrain, including the mesencephalic dopaminergic (mDA) neurons of the ventral tegmental area (VTA) (3). These neurons innervate several cerebral structures regulating complex behaviors (10,11) and require *Otx2* expression for their proper development and survival (12,13), suggesting implication in psychiatric disorders.

A role for Otx2 in juvenile and adult control of complex traits such as depression or anxiety has been revealed by maternal separation experiments. A recent study showed periodic separation of mouse pups from their mothers, during a critical period from postnatal day 10 (P10) to P20, results in a transient decrease in *Otx2* expression in the VTA that leads directly to latent anxiety and depressive phenotypes in male adults (14). In a separate study of anxiety induced by maternal separation, it was proposed that *Otx2* upregulation in the ChP and increased OTX2 levels in the ventral hippocampus inhibitory interneurons may participate in the anxiety phenotype (15). While the first study implicates cell-autonomous activities for OTX2, this second study suggests non-cell-autonomous functions.

We have shown that OTX2 is secreted by ChP into the cerebrospinal fluid (CSF) and transported to the brain parenchyma where it is primarily captured by the fast-spiking GABAergic interneurons that express parvalbumin (PV), hereafter referred to as PV cells (16). The specificity of this capture is mediated by a glycosaminoglycan-binding motif in OTX2 that results in an interaction with proteoglycan-rich perineuronal nets (PNNs) that enwrap cortical PV cells (17,18). During mouse postnatal development, the privileged accumulation of OTX2 within PV cells has been shown to regulate critical periods of plasticity in the primary visual cortex, primary auditory cortex and medial prefrontal cortex (mPFC) (19,20). Furthermore, pharmacological or genetically-induced decrease of OTX2 content in PV cells of the adult cortex reopens a window of plasticity, allowing for the recovery of binocular vision in a mouse model of amblyopia (16,17).

It has been hypothesized that critical periods impact the development of mood disorders such as anxiety and schizophrenia (21–23). Supporting the above-mentioned maternal separation results, we recently found that the *Otx2^+/AA^* mice display prolonged acquisition of acoustic preference in adulthood linked to anxiolysis (20). This constitutive mouse model shows reduced affinity for PNNs, which results in decreased accumulation of OTX2 in PV cells and delayed critical period timing across modalities, suggesting that this transfer may regulate complex animal behaviors. However, it is not clear whether this regulation of a complex behavior is due to purely non-cell-autonomous CSF-borne OTX2 activity or includes some contribution by cell-autonomous OTX2 function in ChP, VTA or other brain region. To begin addressing this important question, we aimed to better characterize the constitutive *Otx2* heterozygote (*Otx2-het*) mouse line (24,25). By using a battery of behavioral tests, we find these mice display anxiolysis-like behavior with no changes in motor activities, cognition, depression-like behavior or sensorimotor gating. This phenotype is recapitulated in mouse models of non-cell-autonomous OTX2 loss-of-function and can be rescued by *Otx2* overexpression in the ChP.

## MATERIAL AND METHODS

### Ethics statement

All animal housing and experimental procedures were carried out in accordance with the recommendations of the European Economic Community (2010/63/UE) and the French National Committee (2013/118). For surgical procedures, animals were anesthetized with Xylazine (Rompun 2%, 5 mg/kg) and Ketamine (Imalgene 500, 80 mg/kg) by intraperitoneal injection. This research (project no. 00704.02) was approved by Ethics committee n° 59 of the French Ministry for Research and Higher Education.

### Mice

The *Otx2-het* mouse line was generated in the laboratory of Antonio Simeone (CEINGE, Naples), with the *Otx2* coding sequence and introns replaced by *GFP* (24). *Otx2^+/GFP^* males were crossed with B6D2F1 females to obtain *Otx2-het* mice. The *Otx2^+/AA^* mouse line was generated through a knock-in approach, as described previously (25). The conditional secreted single-chain antibody (scFv) OTX2 *scFvOtx2^tg/0^* mouse line was generated by targeted transgenics in the *Rosa26* locus, as described previously (26). Mice were raised in a 12-hour light/dark cycle with 2 to 5 (male) or 6 (female) animals per cages. Temperature was controlled (21±2°C), and food and water provided ad libitum.

### Stereotaxic injections

Adeno-associated viruses (AAV) were generated by Vector Biolabs: AAV5(CMV)HAOtx2-2A-mCherry and AAV5(CMV)mCherry. Bilateral intracerebroventricular (icv) stereotaxic injections (bregma: x = −0.58 mm, y = ±1.28 mm, z = 2 mm) of 2 μl high-titer AAV (~10^12^ GU/ml) were performed with a Hamilton syringe at a rate of 0.3 μl/min. Mice were used for biochemical, histological and behavioral analysis at least 3 weeks after infection. Cre-TAT icv injections were performed in *scFvOtx2^tg/0^* mice as previously described (16). Behavior was tested 4 weeks after injection.

### Behavior analyses

Tests were performed from P90 to P120 in littermate groups of female or male separately in the following order: locomotor activity, Y-maze, elevated plus maze, light-dark box, rotarod, tail-suspension, prepulse inhibition, forced swimming test. Except for rotarod, tail-suspension and prepulse inhibition tests, behavior was registered using a semi-automated infrared system (Viewpoint, Lyon). All tests were done during the light cycle between 9:00 and 18:00. Animals were habituated to the testing room for at least 30 min and the experimenter was blind to animal genotype or treatment. To eliminate odor cues, each apparatus was thoroughly cleaned after each animal.

#### Y maze

Short-term spatial memory and locomotor activity was evaluated by spontaneous alternative exploration of the Y-maze during 10 min. The maze consisted of 3 alleys diverging at 120° from one another, 40 cm long, 15 cm width, with black 30 cm high Perspex borders decorated differently in each arm, which were illuminated at 50 lux. The mouse was placed at the tip of one arm and the time spent visiting each of the other arms sequentially was counted as a memory performance.

#### Elevated plus maze

A cross-shape maze elevated at 70 cm from the floor was illuminated at 50 lux. Two “closed” arms facing each other and protected by black Perspex walls (20 cm high), are crossed perpendicularly by the two “opened” unprotected arms. The mouse was placed at the center of the cross and activity was recorded for 10 min. Total distance traveled, the time spent in the open arms and the number of entries into each arm were measured.

#### Light-dark box

Boxes are composed of a dark (black walls, floor and roof) and light (white walls and floor, no roof, 150 lux) compartments of equal volume (20 x 20 x 30 cm) linked by a small opening. The mouse is placed in the light box facing the opening, and anxiety is measured by the time spent in the light compartment over a 10 min period.

#### Rotarod

Equilibrium and motor coordination were measured in a progressively accelerating (from 2 to 40 rpm during 5 min) rotarod (model 47600, Ugo Basile, Italy). Three daily trials separated by 90 min intervals were performed during 2 days.

#### Tail-suspension test

Mice were suspended by the tail with adhesive linked to a gauge detecting movement (automated system BIOSEB, France), during 6 min. Immobility was interpreted a sign of depression-like behavior.

#### Forced swimming test

Depressive behavior was evaluated by registering the length of freezing periods of mice placed in a see-through Perspex cylinder (h, 25 cm; d, 13 cm) full of water (h, 20 cm; 25 °C). On the first day, mice were habituated during 10 min; mice were tested on the second day during a period of 6 min. An infrared panel positioned vertically behind the cylinder detects mice movements. Immobility was interpreted a sign of depression-like behavior.

#### Prepulse Inhibition test

Testing was carried out in a SR-Lab system (San Diego Instruments, USA). Each mouse was placed in a startle chamber, and acoustic noise bursts were presented via a speaker mounted 26 cm above. Throughout the session, a background noise level of 68 dB was maintained. After a 5-min acclimatization period (68 dB background noise), 10 startle pulses (120 dB, 40 ms duration) were presented with an average inter-trial interval of 15 s. During the next 20 min, no stimulus (background noise, 68 dB), prepulses alone (72, 76, 80 or 84 dB, 20 ms duration), startle pulses alone, and prepulses followed 80 ms later by startle pulses were presented 10 times, randomly distributed. Percent prepulse inhibition (PPI) was calculated as [100 – 100x (startle response of acoustic startle from acoustic prepulse and startle stimulus trials)/ (startle response alone trials)].

### Dosage of amines and amine metabolites

Brains were dissected in ice-cold PBS and dry tissue was kept at −80°C. Extracts were prepared and analyzed by HPLC using a reverse phase column (Nucleosil 120-3 C18; Knauer) with electrochemical detection as described (27). Data were recorded and quantified using HPLC Chromeleon computer system (Dionex).

### Quantitative RT-PCR

Mice were sacrificed by cervical dislocation, and mPFC, amygdala, hippocampus and/or ChP were microdissected in ice-cold PBS and frozen in liquid nitrogen. Total RNA was extracted with the AllPrep DNA/RNA Mini Kit (Qiagen 80204), with on-column genomic DNA digestion, and evaluated by spectroscopy (Nanodrop, Palaiseau, France). cDNA was synthesized from 500 ng of total RNA with QuantiTect Reverse Transcription kit (Qiagen 205313). Quantitative PCR reactions were carried out in triplicate with SYBR Green I Master Mix (Roche S-7563) on a LightCycler 480 system (Roche). Expression was calculated by using the 2^-ΔΔCt^ method with *Ywhaz* as a housekeeping reference gene. Primer sequences: *PV*, Fwd-AAGAAACAAAGACGCTTCTGGC, Rev-ACTGAACAGAAACTCAGGAGGG; *SST*, Fwd-CTGCGACTAGACTGACCCAC, Rev-AAAGCCAGGACGATGCAGAG; *Gadd45b*, Fwd-TCTCTAGAGGAACGCTGAGACC, Rev-GTAGGGTAGCCTTTGAGGGATT; *Arc*, Fwd-AGAGCTGAAGGTGAAGACAAGC, Rev-CAAGAGGACCAAGGGTACAGAC; *Nr4a1*, Fwd-AGGAGACCAAGACCTGTTGCTA, Rev-TGTAGTACCAGGCCTGAGCAGA; *mt-Nd4*, Fwd-AATATACATAATTATTACCACCCAACG, Rev-TGTCAGACCTGTAATTAGTTTTGGA; *Iba1*, Fwd-CTCAGCTCACCCCATTCCTG, Rev-ACATCAGCTTCTGTTGAAATCTCC; *Ywhaz*, Fwd-TTGATCCCCAATGCTTCGC, Rev-CAGCAACCTCGGCCAAGT.

### Immunohistochemistry

Anesthetized mice were perfused transcardially (10 ml/min) with 20 ml of PBS and 30 ml of 4% paraformaldehyde in PBS. Dissected brains were post-fixed in 4% paraformaldehyde at 4 °C for 1 h, rinsed in PBS, soaked in 20% sucrose-PBS at 4°C for 12-24 h and frozen. Fluorescent immunohistochemistry was performed on cryostat sections (20 μm). Briefly, after permeabilization with 1% Triton for 20 min, sections were incubated in 100 mM glycine for 20 min, blocked with PBS, 0.2% Triton, 10% normal goat serum (NGS) for 45 min and incubated overnight with primary antibodies: anti-tyrosine hydroxylase antibody (rabbit, abcam, 1/500); anti-dopamine transporter (rat, Millipore, 1/5000); biotinylated *Wisteria floribunda* agglutinin (1/100, Sigma L1516); anti-parvalbumin (rabbit, Swant, 1/500); anti-tryptophan hydroxylase 2 (rabbit, abcam, 1/500); anti-OTX2 (mouse monoclonal, in house) in PBS, 1% Triton, 3% NGS at 4 °C. After 3 washes in PBS, 1% Triton, secondary antibodies were incubated (Alexa Fluor-conjugated, Molecular Probes) for 2 h at room temperature (1/2000). After 3 washes in PBS, 1% Triton, sections were mounted in DAPI-fluoromount medium and kept at 4 °C.

DAB-immunohistochemistry was performed on free-floating cryostat sections (40 μm). Endogenous peroxidase was removed with a 5 min incubation in 0.3% H2O2, 0.2% Methanol. Sections were permeabilized and blocked as above and incubated during 12-24h with an anti-tyrosine hydroxylase antibody (rabbit, Pel Freeze Biologicals, 1/1000) in PBS, 0.1% Triton, 3% NGS at 4°C. After 3 washes in PBS, 0.1% Triton, a biotin-coupled anti-rabbit antibody (1/2000) was added for 2h at room temperature. After 3 washes in PBS, 0.1% Triton, biotin was detected by using the Vectastain Elite ABC HRP kit and Peroxidase Substrate kit (1h of ABC buffer + reaction with DAB, Vectorbiolabs), and the slides were washed in water prior to mounting.

Images were acquired with either a Leica SP5 or SP8 confocal microscope. For VTA cell counting, every second section (containing VTA) was counted by stereology using Stereo Investigator software (MBF Bioscience Inc.) at a 40X magnification. Amygdala were imaged at 20X and the density of fiber staining was assessed by the mean grey value option with ImageJ software. The infralimbic mPFC were imaged at 20X and 40X and cell number and staining intensity was measured with ImageJ and in-house macros.

### Statistical Analysis

All statistics were performed using Prism (version 8.1.2, GraphPad Software). In order to avoid type II errors, outliers were defined as more than 2 standard deviations from the group mean and were removed from analysis. Main effects and interactions with more than 2 groups were determined using analyses of variance (ANOVA) or 2-way ANOVA with Tukey’s multiple comparison *post hoc* test. The *t*-test was used for single comparisons among two groups.

## RESULTS

### *Otx2-het* mice show anxiolysis-like behaviors

Three-month-old *Otx2-het* and wild-type (WT) littermates were analyzed for motor coordination (rotarod), depression-like behavior (tail-suspension and forced swimming tests), cognition (Y-maze), sensorimotor gating (prepulse inhibition) and anxiety (light-dark box and elevated plus maze). Motor coordination measured by the rotarod test showed differences between males and females (F(3, 270) = 28.50; *p* < 0.0001), but not between WT and *Otx2-het* animals (Supplementary Figure 1A). Depression-like behavior measured by the forced swimming (Supplementary Figure 1B) and tail suspension (Supplementary Figure 1C) tests showed no differences between the two genotypes and a small difference between males and females in the tail suspension test only (77.09 ± 12.84 vs 39.09 ± 5.41; F (1, 52) = 10.40; *p* = 0.0022), with females showing less effort to escape. The short-term spatial memory test (Y maze) was used to measure both locomotor activity and memory. Although females show a slightly higher motor activity (4944 ± 130 vs 4439 ± 160; F(1, 54) = 4.75; *p* = 0.0337), here again no differences were found between the two phenotypes (Supplementary Figure 1D). Another absence of difference between genotypes was found in the prepulse inhibition (PPI) test (Supplementary Figure 1E, F). These results suggest the loss of one *Otx2* allele does not affect motor activities, cognition, depression-like behavior or sensorimotor gating.

In contrast, strong differences between WT and *Otx2-het* mice appeared in the light-dark box (LDB) and the elevated plus maze (EPM) anxiety tests (Figure 1A, B). *Otx2-het* mice spent more time than WT in the light compartment of the LDB (58.84 ± 9.64 (WT) vs 115.93 ± 13.87; F(1, 57) = 11.34; *p* = 0.0014) (Figure 1A) as well as in the open arms of the EPM (6.00 ± 1.44 (WT) vs 46.13 ± 8.71; F (1, 52) = 18.49; *p* < 0.0001) and with increased percentage of entries in the open arms (6.83 ± 1.29 (WT) vs 16.08 ± 2.27; F(1, 58) = 13.35; *p* = 0.0006). Furthermore, the increases in the open arms visits and time are in parallel with an increase of the distance travelled by the animals in the EPM (2.21 m ± 0.11 (WT) vs 2.68 cm ± 0.16; F(1, 56) = 6.05; *p* = 0.017). These effects are similar in both sexes given the absence of significant sex-genotype effect in each test (Figure 1). The EPM test was also conducted in absence of light (Figure 1C) ensuring that the anxiolysis-like behavior associated with the loss of one *Otx2* allele is independent of a putative vision deficit created by decreased *Otx2* expression (25) (6.86 ± 1.78 (WT) vs 25.34 ± 5.67; F(1, 42) = 10.09; *p* = 0.0028), showing again increased percentage of entries in the open arms in the mutants compared with the WT (6.44 ± 3.15 (WT) vs 14.03 ± 0.47; F(1, 44) = 7.07; *p* = 0.011) and the total distance travelled by the mutant mice (2.02 m ± 0.17 (WT) vs 2.70 m ± 0.26; F(1, 43) = 4.74; *p* = 0.035). The EPM test was thereafter used as an anxiety read-out in all experiments.

**Figure 1.**
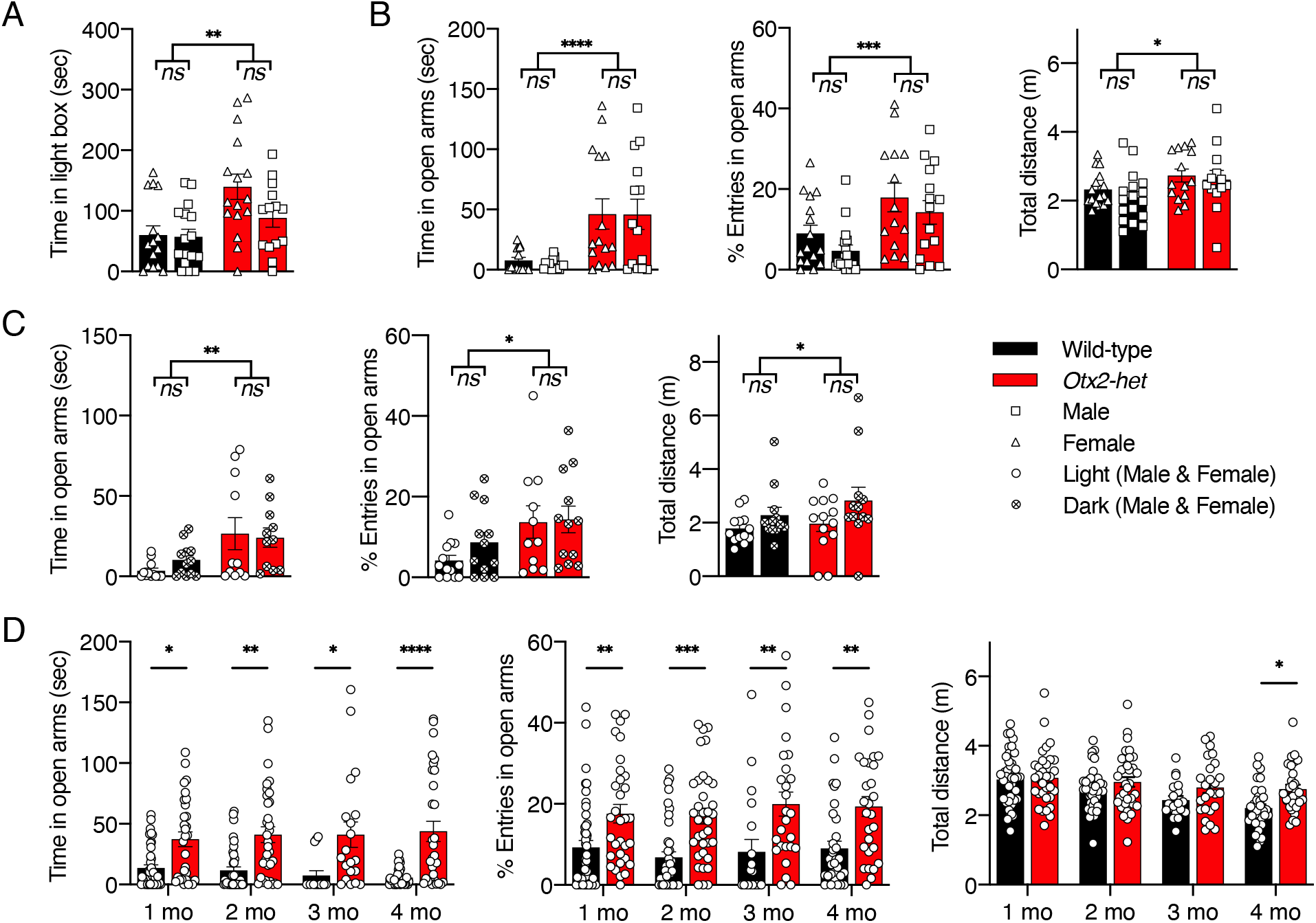
Sex indifferent anxiolysis-like behavior in *Otx2-het* mice. **(A)** Light-dark box test of adult mice (P90 to P120) during a 10 min period (WT females n=15; WT males n=16; *Otx2-het* females n=16; *Otx2-het* males n=14). **(B)** Elevated plus maze test during a 10 min period (WT females n=15; WT males n=15; *Otx2-het* females n=15; *Otx2-het* males n=14). **(C)** Elevated plus maze performed in normal (50 lux) or dark (0 lux) conditions (WT light n=13; WT dark n=12; *Otx2-het* light n=11; *Otx2-het* dark n=12). **(D)** Elevated plus maze test at different ages (WT P30 n=44; WT P60 n=39; WT P90 n=16; WT P120 n=33; *Otx2-het* P30 n=32; *Otx2-het* P60, n=35; *Otx2-het* P90 n=22; *Otx2-het* P120 n=31). All values: mean ± SEM; two-way ANOVA, post hoc Tukey test; \**p* < 0.05, \*\**p* < 0.01, \*\*\**p* < 0.001, \*\*\*\**p* < 0.0001.

### The low anxiety behavior of *Otx2-het* mice in the EPM test does not reflect a dopaminergic or serotoninergic phenotype

In a previous study, it was found that mice heterozygous for *Engrailed-1*, a homeogene expressed in the mDA neurons of the ventral midbrain, experience a progressive loss of these neurons in the substantia nigra (SNpc) and VTA beginning at 6 weeks (28). This results in a 40% and 20% neuronal loss at 48 weeks in the SNpc and VTA, respectively. *Otx2* is expressed in the adult VTA (13), and the projections of the VTA into several target structures, including the hippocampus, nucleus accumbens, amygdala, and the mPFC, may be involved in anxiety regulation (14). This led us to analyze the hypoanxiety behavior of *Otx2-het* mice between 1 and 4 months of age, including the age at which behavior was tested (3 months) to determine whether the behavior was appearing/disappearing with the age (Figure 1D). We observed no changes in the anxiety level in any of the interrogated time points when time in open arms was evaluated (F(1, 229) = 56.06;*p* < 0.0001; 1 mo,*p* = 0.03; 2 mo, *p* = 0.0019; 3 mo,*p* = 0.032; 4 mo, *p* < 0.0001) and percentage of entries in the open arms (F(1, 248) = 45.45; *p* < 0.0001; 1 mo, *p* = 0.098; 2 mo, *p* = 0.001; 3 mo, *p* = 0.0038; 4 mo, *p* = 0.0028). For the distance traveled by the animals we only observed differences at 4 months of age (F (1, 248) = 10.24; *p* = 10.24; 4 mo, *p* = 0.0016). Thus, anxiolysis-like behavior is established very early and is maintained during this period, making it unlikely that it is due to mDA progressive neuronal loss.

This hypothesis was further confirmed by stereology counting of tyrosine hydroxylase (TH)-positive cell bodies of the VTA (Figure 2A), which showed no differences between 3-month-old *Otx2-het* mice and their WT littermates. However, as illustrated in a study on SNpc mDA neurons in WT and *Engrailed-1* heterozygotes, cells bodies can be present while axons begin degenerating as characterized by their fragmentation, clearly visible both in the medial forebrain bundle and in the striatum (29). Hence, we evaluated the state and density of the TH- and dopamine transporter (DAT)-positive fibers in the basolateral amygdala, a region responsible for the fear response and receiving mDA terminals from the VTA (30). No significant changes were observed neither for the density of any of the two markers of dopaminergic neurotransmission, nor for the morphology of the fibers (Figure 2B).

**Figure 2.**
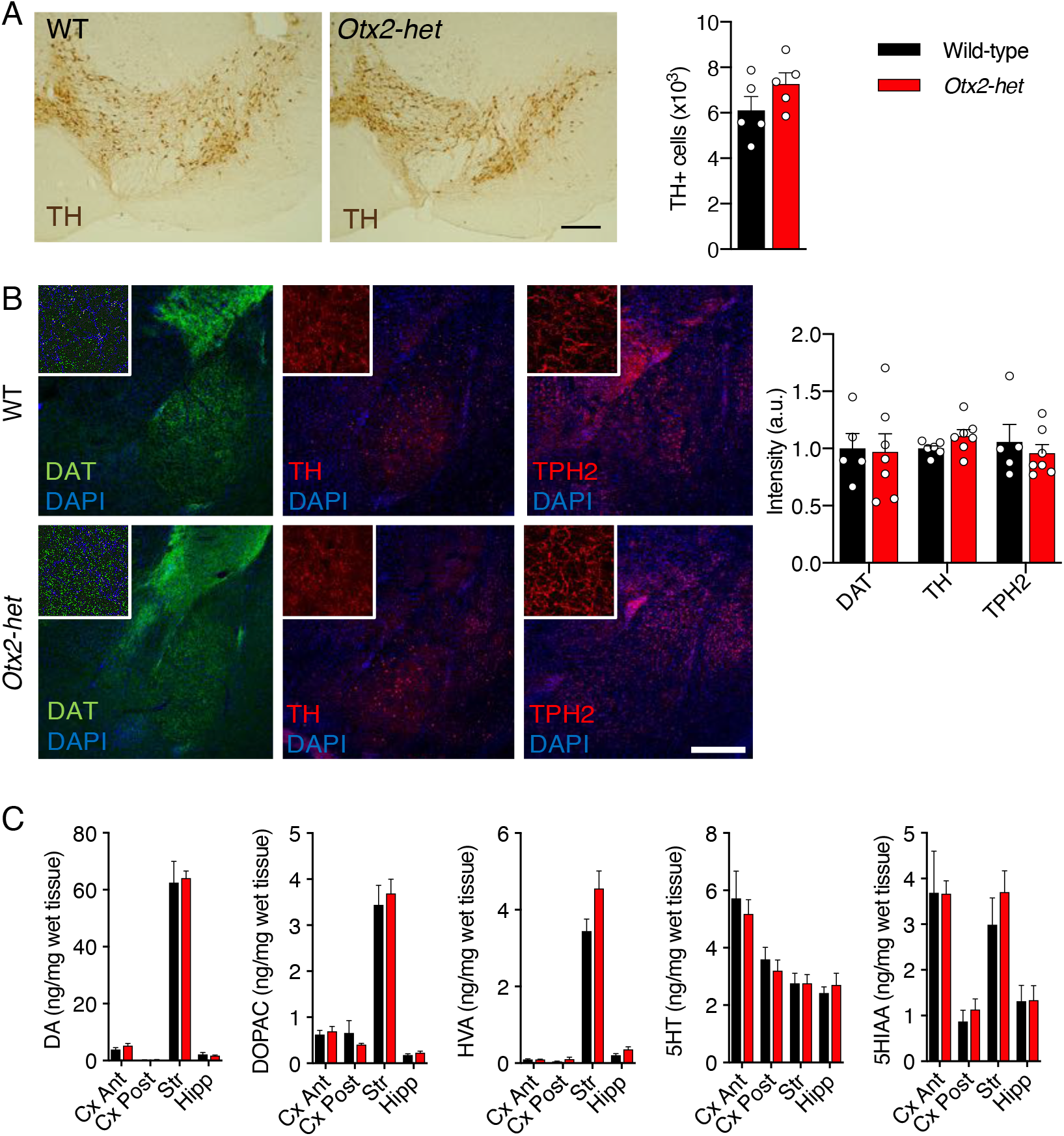
Dopaminergic and serotoninergic markers are not disturbed in the *Otx2-het* mice. **(A)** Representative images and quantification of tyrosine hydroxylase (TH) immunostaining in the ventral tegmental area (VTA) of *Otx2-het* (n=4) and WT (n=5) mice at ~P90 (scale bar: 50 μm). **(B)** Quantification of metabolites related to the dopaminergic and serotoninergic systems in extracts from anterior cortex (Cx Ant) and posterior cortex (Cx Post), striatum (Str), and hippocampus (Hipp) of ~P90 mice (WT n=5; *Otx2-het* n=7). DA, dopamine; DOPAC, 3,4-dihydroxyphenylacetic acid; HVA, homovanillic acid; 5HT, 5-hydroxytryptamine; 5HIAA, 5-hydroxyindoleacetic acid. **(C)** Representative images and quantification of immunostaining of fibers for dopamine transporter (DAT), TH, and tryptophan hydroxylase 2 (TPH2) in the basolateral amygdala (BLA) of *Otx2-het* (n=7) and WT (n=6) mice (scale bar: 250 μm). All values: mean ± SEM; *t*-test.

To further evaluate a possible dopaminergic phenotype, dopamine (DA) and its main metabolites dihydroxyphenylacetic acid (DOPAC) and homovanillic acid (HVA) were measured in the cortex (anterior and posterior), the striatum, and the hippocampus of 3-month-old WT and *Otx2-het* mice from the same litter (Figure 2C). The amounts are very similar in both genotypes and in all studied structures, thus eliminating both a change in absolute DA concentrations and in DA metabolism.

Early *Otx2* hypo-morphism can also lead to an increase in the number serotonin (5HT) neurons by an anterior shift in the midbrain/hindbrain boundary (31,32). Because 5HT has been implicated in anxiety regulation (33,34), we measured the levels of 5HT and of its main metabolite 5HIAA. We found no differences between animals of either genotype and in any of the studied structures (Figure 2C). Furthermore, the serotoninergic fibers in the amygdala marked with the TPH2 antibody showed similar morphology and density in both WT and *Otx2*-het mice (Figure 2B).

### Non-cell-autonomous OTX2 mouse models recapitulate anxiolysis-like behavior

The absence of VTA-dependent changes led us to consider non-cell-autonomous OTX2 activity. Indeed, previous analysis of the *Otx2^+/AA^* mouse model, which results in reduced OTX2 transfer to cortical PV cells, revealed a music preference phenotype linked to anxiolysis (20). We confirmed anxiolysis-like behavior in *Otx2^A^* mice in the EPM tests (Figure 3A). Both male and female mice showed increased time in open arms (15.24 ± 2.28 (WT) vs 25.64 ± 3.08; F(1, 49) = 4.1; *p* = 0.048) and higher number of entries (8.27 ± 1.34 (WT) vs 14.81 ± 0.71; F(1, 48) = 7.86; *p* = 0.0073), with no change in total distance. However, no significant differences were observed for the LDB test (Figure 3B).

**Figure 3.**
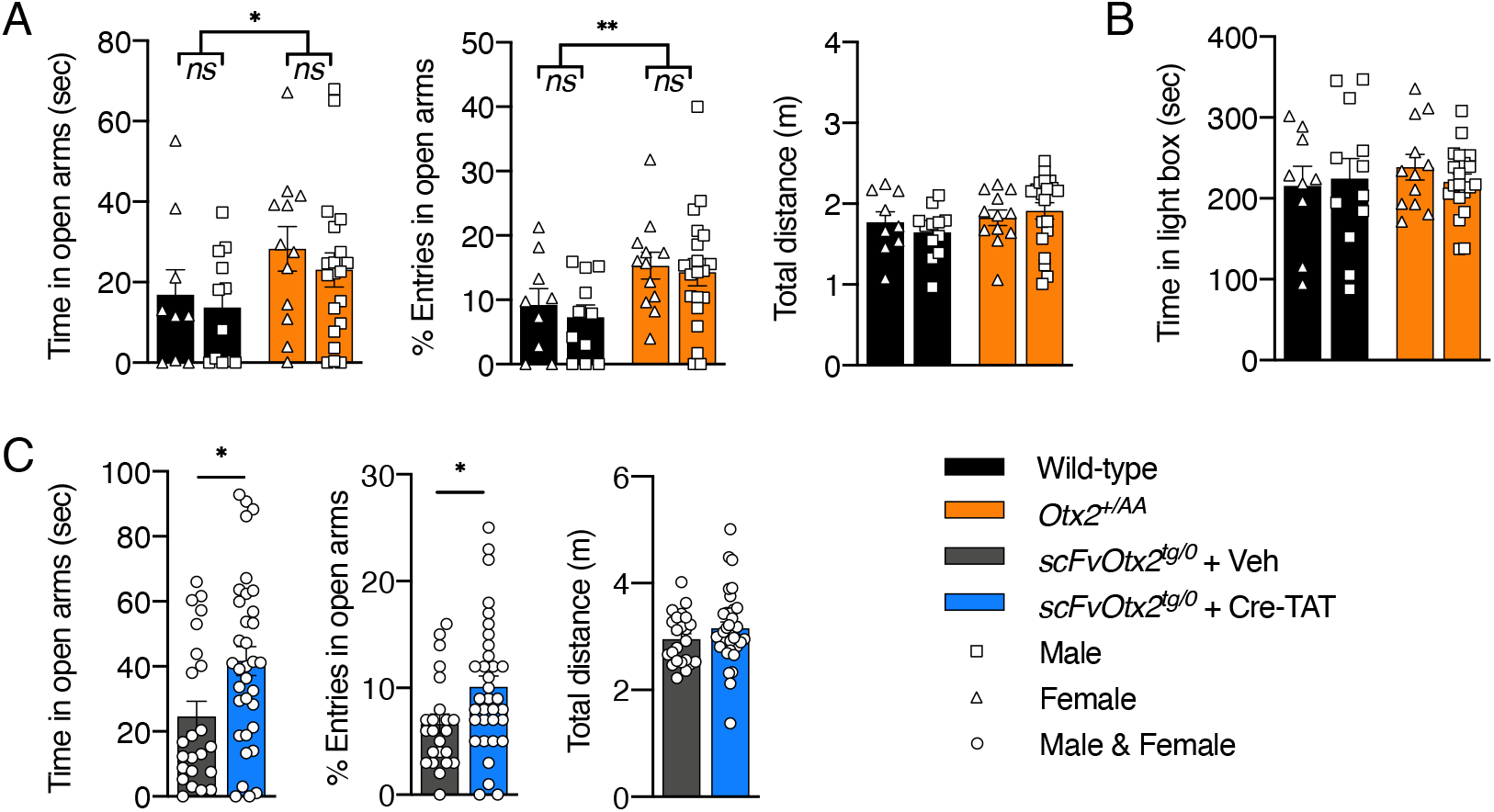
Anxiolysis-like behavior in non-cell autonomous OTX2 knock-down mouse models. **(A)** Elevated plus maze test of *Otx2^+/AA^* mice at P90 during a 10 min period (WT females n=9; WT males n=12; *Otx2^+/AA^* females n=12; *Otx2^+/AA^* males n=20). **(B)** Light-dark box test of *Otx2^+/AA^* mice at P90 during a 10 min period (WT females n=15; WT males n=16; *Otx2^+/AA^* females n=16; *Otx2^+/AA^* males n=14). **(C)** Elevated plus maze test of *scFvOtx2^tg/0^* mice at ~P100, 15 days after intracerebroventricular (icv) injection of vehicle (Veh, n=23) or Cre recombinase (Cre-TAT, n=34). All values: mean ± SEM; two-way ANOVA, post hoc Tukey test in **A** and **B**; *t*-test in **C**; \**p* < 0.05, \*\**p* < 0.01.

The observed changes in critical period timing of the *Otx2^+/AA^* mouse model were hypothesized to be due to altered distribution of CSF-borne OTX2 into mPFC PV cells, resulting in accompanying changes in PV and PNN staining and alteration in mPFC circuit recruitment (20). The ChP not only cleans the CSF from toxic metabolites but also nurtures the brain by secreting into the CSF a large number of trophic factors (35,36), including OTX2 (16). To verify if OTX2 secreted by the ChP could regulate anxiety, we took advantage of a conditional mouse line expressing a secreted scFv directed against OTX2 (26). In this mouse, injection of a cell-permeable Cre recombinase in the lateral ventricles of the brain induces the secretion of the scFv-OTX2 antibody from the ChP, which leads to the neutralization of extracellular OTX2 in the CSF and, in the case of the visual cortex, to the reopening of plasticity in the adult mouse. Through EPM tests, we find that neutralizing ChP-derived OTX2 in the CSF of *scFvOtx2^tg/0^* mice installs an anxiolysis-like phenotype at P90 (Figure 3C). These mice showed increased time in open arms (24.60 ± 4.66 (WT) vs 41.69 ± 4.47; *t* = 2.568; *p* = 0.0130) and higher number of entries (6.65 ± 0.89 (WT) vs 10.09 ± 1.05; *t* = 2.328; *p* = 0.0236), with no change in total distance.

### PV expression is reduced in the mPFC of *Otx2-het* mice

The anxiety response is tightly regulated by the mPFC (37), which is mediated in part by PV cells (38). The above results strongly suggest that non-cell autonomous OTX2 plays a role in mouse anxiety phenotypes. In postnatal mice, *Otx2* is expressed in several extra-cortical structures, including the VTA and ChP (14,16) but no expression takes place in the cerebral cortex parenchyma (19). Although not expressed in the cerebral cortex, OTX2 is imported from extra-cortical structures, in particular ChP, by neurons throughout the cortex (primarily PV cells), including primary cortices and limbic structures. Non-cell autonomous OTX2 has been identified in mPFC, hippocampus, and basolateral amygdala (15,16,20). Because of this presence of OTX2 in several structures implicated in the regulation of anxiety-related behaviors, we tested the expression of different genes in these regions to identify possible altered pathways due to potentially reduced OTX2 level (Figure 4A). We quantified mRNA levels corresponding to interneuron marker genes (*PV* and *SST*), plasticity genes known to be altered by OTX2 levels (*Gadd45b, Arc, Nr4a1*), and inflammatory/oxidative stress pathway genes (*Mtnd4* and *Iba1*), given that mood disorders can be related to oxidative stress response (39,40). While no significant changes were observed in the amygdala or the hippocampus, PV gene expression was significantly reduced in the mPFC of *Otx2-het* mice (*t* = 2.293; *p* = 0.04). This change in PV expression was confirmed by the decreases number of cells stained for PV in the infralimbic mPFC of *Otx2-het* mice at P90 (Figure 4B, *t* = 2.873; *p* = 0.0263). Furthermore, the number mature PV cells, as defined by PNN intensity, is also compromised; PV+WFA+ cell number is significantly reduced (*t* = 3.180; *p* = 0.0107), along with overall WFA staining intensity per cell (*t* = 2.773; *p* = 0.0216). These results suggest that the observed anxiolysis-like phenotypes are due to changes in infralimbic mPFC PV cell populations.

**Figure 4.**
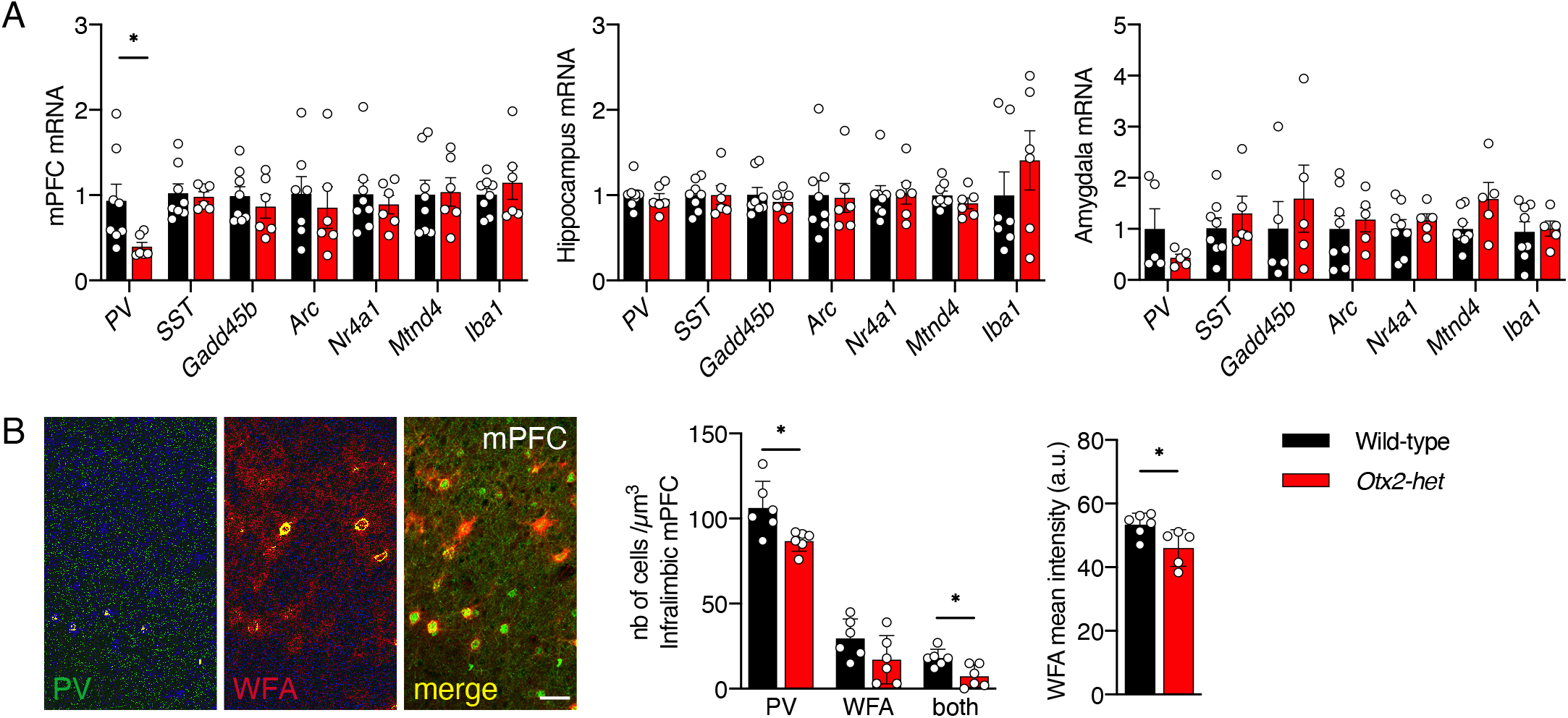
Downregulated parvalbumin (PV) expression in the medial prefrontal cortex (mPFC) of *Otx2-het* mice. **(A)** Analysis of mRNA levels of genes associated with interneurons (*PV*; *SST*, somatostatin), plasticity (*Gadd45b*, Growth arrest and DNA-damage-inducible beta; *Arc*, activity-regulated cytoskeleton-associated protein; *Nr4a1*, nuclear receptor subfamily 4 group A member 1), and metabolism/inflammation (*Mtnd4*, mitochondrial NADH-ubiquinone oxidoreductase chain 4; *Iba1*, ionized calcium-binding adapter molecule 1) in extracts from the mPFC, hippocampus, and amygdala of WT (n=8) and *Otx2-het* (n=6) mice at ~P90. Data are normalized to *Ywhaz* levels of WT mice. **(B)** Representative images and quantification of PV and *Wisteria floribunda* agglutinin (WFA) staining in the infralimbic mPFC of WT (n=6) and *Otx2-het* (n=6) mice at P90 (scale bar: 50 μm). All values: mean ± SEM; *t*-test; *p < 0.05.

### OTX2 secreted by the ChP regulates mouse anxiety

To evaluate the role of ChP-derived OTX2 in the *Otx2-het* mice anxiety levels, we performed a rescue experiment involving ivc injection of AAV serotype 5 encoding *Otx2*. This serotype provides specific infection of the ChP (Figure 5A) and resulted in ~1.5-fold increase in *Otx2* expression (Figure 5B, *t* = 3.525; *p* = 0.0023). This overexpression caused increased anxiety in the *Otx2-het* mice in the EPM test (Figure 5C), resulting in decreased time in the open arms (15.11 ± 3.58 (*Otx2-het*) vs. 6.01 ± 1.54; *t* = 2.286; *p* = 0.029) and less entries (19.56 ± 2.02 (*Otx2-het*) vs. 12.78 ± 2.24; *t* = 2.252; *p* = 0.031), but with no change in the total distance traveled. Histochemical analysis of PV cell maturation (Figure 5D) showed a trending increase in PV cell number (*t* = 1.925; *p* = 0.0864) with a significant increase in PNN+ PV cells (*t* = 3.173; *p* = 0.0113) and WFA staining intensity (*t* = 2.773; *p* = 0.0216). These experiments suggest that OTX2 expression in the ChP is sufficient to restore a normal anxiety behavior in the *Otx2-het* mouse through accompanying changes in mPFC PV cell populations.

**Figure 5.**
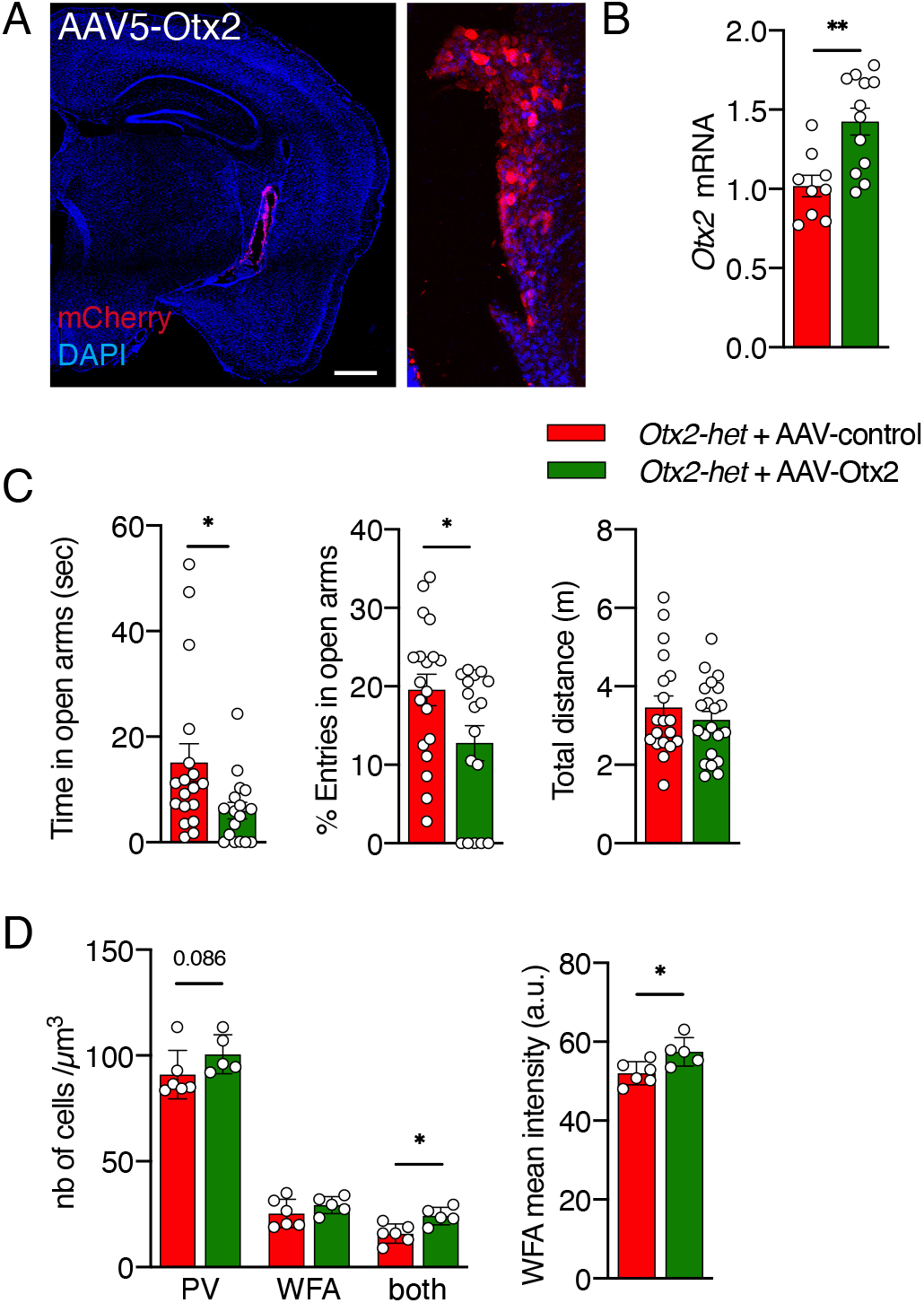
Overexpression of OTX2 in the choroid plexus *Otx2-het* mice increases anxiety. **(A)** Image of mCherry staining in a coronal section of a P100 *Otx2-het* mouse, 3 weeks after intracerebroventricular (icv) injection of AAV5(CMV)HAOtx2-2A-mCherry virus, which specifically infects the choroid plexus (scale bar: 100 μm). **(B)** Analysis of *Otx2* mRNA levels (normalized to *Ywhaz*) in choroid plexus extracts 3 weeks after icv virus injection (AAV5-control n=9; AAV5-Otx2 n=12). **(C)** Elevated plus maze test of *Otx2-het* mice during a 10 min period, 4 weeks after icv injection of AAV5-control (n=19) or AAV5-Otx2 (n=17). **(D)** Quantification of parvalbumin (PV) and *Wisteria floribunda* agglutinin (WFA) staining in the infralimbic medial prefrontal cortex (mPFC) of *Otx2-het* mice, 4 weeks after icv injection of AAV5-control (n=6) or AAV5-Otx2 (n=5). All values: mean ± SEM; *t*-test; *p < 0.05, **p < 0.01.

## DISCUSSION

In this study, we provide evidence for a role of Otx2 in the regulation of anxious behavior and show that OTX2 secreted from the ChP into the CSF plays a role in this regulation. This does not exclude other extra-cortical sources, in particular the pineal gland, but establishes that the ChP, through the synthesis and secretion of OTX2 is an important anxiety regulator. Anxiety regulation can be the consequence of modest changes in OTX2 expression and/or secretion. For expression, the AAV5 strategy induces a 50% increase in ChP *Otx2* transcription. For secretion, previous studies using the same neutralization strategy in the CSF showed that a 20% OTX2 decrease in the visual cortex PV cells is sufficient to reopen plasticity in the adult (26). In the present study, all target structures were not identified, but it can be claimed that an acute decrease of OTX2 en route from the ChP to the brain parenchyma of adult mice (~P90) is enough to decrease anxiety. This result excludes that *Otx2-het* mice anxiolysis-like behavior is purely developmental, as also suggested by the similar reduced anxiety of *Otx2-het* mice between 1 and 4 months of age.

Interestingly, the *Otx2^+/AA^* mutant showed anxiolysis-like behavior in the EPM but not the LDB tests. According to analysis of retina structure and function, this mutant shows OTX2 activity level ~75% of normal, while the *Otx2-het* mouse shows ~50% compared to WT (25). Furthermore, the *Otx2^+/AA^* mouse shows delays in critical period timing that are not as extended as in the *Otx2-het* mouse (4,19,20), suggesting a subtler phenotype is to be expected. Nevertheless, the EPM test appears to more sensitive than the LDB test for measuring anxiety.

While all non-cell autonomous OTX2 target cells and structures were not investigated, in such complex behaviors, it is likely that several interacting structures are implicated. If so, the presence of OTX2 in the CSF could give the protein access to many candidate target structures functioning in association through the existence of neuronal networks and hub structures (41,42). This is apparently the case as non-cell autonomous OTX2 was found in PV cells from the mPFC, ventral hippocampus and amygdala (15,16). Interestingly, recent experiments strongly suggest that OTX2 imported in PNN-enwrapped PV cells in the mPFC and in the ventral hippocampus might regulate anxiety (15,20). Although the accent here is placed on PV cells, other interneurons might play an OTX2-associated role as suggested in the mPFC where OTX2 is mainly imported by PV+ and calretinin+ interneurons (70% and 20% co-labeling with OTX2, respectively) (20).

The large repertoire of anxiety-related interconnected target structures is also shared by projections from the VTA (10,11). Since Otx2 is expressed in the adult VTA and may regulate the survival of the mDA cells that constitute 65% of the neuronal population in this structure (43), a possible cell-autonomous role of OTX2 on VTA mDA neuron physiology/survival or, alternatively, a non-cell autonomous effect on their hippocampal of cortical targets following its anterograde transport and trans-synaptic passage cannot not be excluded. The experiments demonstrating that the number of mDA neurons is not affected in the mutant at 3 months of age (the anxiolysis-like phenotype is present already at 1 months and persists at least for 3 more months) is not in favor of the idea that mDA neuron number is at the origin of the phenotype. While we cannot preclude that there were compensatory changes in DA and/or 5HT receptor activity in target regions, we found no change in the amount of DA and DA metabolites in the cortex, hippocampus and striatum, further supporting the view that the mDA “phenotype” is not altered in *Otx2-het* mice. Nor did we find any change in 5HT innervation or the amount of 5HT and of its metabolite 5HIAA which would be expected if the early *Otx2* hypomorphism had shifted the midbrain/hindbrain boundary in a more anterior position (31,32).

The normal number of mDA neurons and the maintained dopaminergic and serotoninergic innervation and metabolism establish that the loss of one *Otx2* allele is not sufficient to significantly modify mDA progenitor numbers, or to significantly decrease mDA neuron survival. However, there are several cases where HP signaling was found to interact with classical signaling pathways. In the fly wing disk, ENGRAILED co-signals with decapentaplegic for the formation of the anterior cross vein (44), while in the chick optic tectum, it interacts with EphrinA5 and adenosine signaling for growth cone guidance or collapse (45,46). A similar interaction was found between PAX6 and netrin signaling in the regulation of oligodendrocyte precursor cell migration (47). In these cases, secreted HP facilitates the full activity of classical signaling. Therefore, we cannot preclude that the anxiolysis-like behavior results from modulation of classical neurotransmitter of growth factor activities by non-cell autonomous OTX2.

Maternal separation experiments have established a role for cell-autonomous OTX2 in the VTA (14), independently of the number of mDA neurons or of DA levels and metabolism. A transient decrease in *Otx2* expression between P10 and P20 in mDA neurons of the VTA can trigger a depressive state in adulthood in male mice following social defeat, suggesting epigenetic alterations independent of cortical plasticity. Our finding that reducing OTX2 in the CSF can reduce anxiety in the adult suggests that the mechanisms involved may differ from those at work in the maternal separation paradigms. Our anxiolysis-like phenotypes across multiple mouse models might be associated with induction of adult cortical plasticity in our conditional models, or with prolonged critical period plasticity in our constitutive models, as previously seen (20). Indeed, another maternal separation paradigm that results in excessive adult anxiety implicates altered ventral hippocampus PV cell plasticity (15), which is accompanied by increased OTX2 in the ChP and non-cell-autonomous OTX2 in the ventral hippocampus. Nevertheless, we cannot rule out that there is a link between *Otx2* expression in the VTA during a critical period (from P10 to P20) and the ability of the ChP to direct OTX2 to non-cell-autonomous target structures.

It has been hypothesized that OTX2 signaling orchestrates complex behaviors reflecting the interplay of multiple sequential critical periods and that the consequence of its disruption is a hallmark of psychiatric and intellectual disorders (3,20). OTX2 regulates PNN and PV expression levels (17), PNN density is found to be low in the prefrontal cortices of schizophrenia patients (48–50), while weakened PV circuits in the mPFC cause deficits in behavioral aspects of schizophrenia patients (51,52). *Otx2* polymorphisms are associated with bipolar disorders (53), and the choroid plexus has been linked with major depressive disorders (54). These findings suggest the potential therapeutic interest of preserving or restoring PV cell function. Our model supports the hypothesis that *Otx2* loss and gain of function in the adult ChP can regulate anxiety levels in the mouse, although it does not preclude that other models may extend the role of *Otx2* expression, during specific developmental periods, to other pathologies. Manipulating OTX2 levels in the ChP and CSF, coupled with behavioral therapies, may hold promising opportunities for psychiatric disorders.

## ACKNOWLEDGEMENTS AND DISCLOSURES

We would like to thank Chantal Dubreuil and Jessica Apulei for technical assistance. Funding was provided by Fondation Bettencourt Schueller, European Research Council Advanced Grant (HOMEOSIGN, ERC-2013-ADG-339379 to A.P.), and Agence Nationale de la Recherche Grant (ANR-11-BLAN-069467 to A.P.). All authors report no biomedical financial interests or potential conflicts of interest.

---

**Figure S1.**
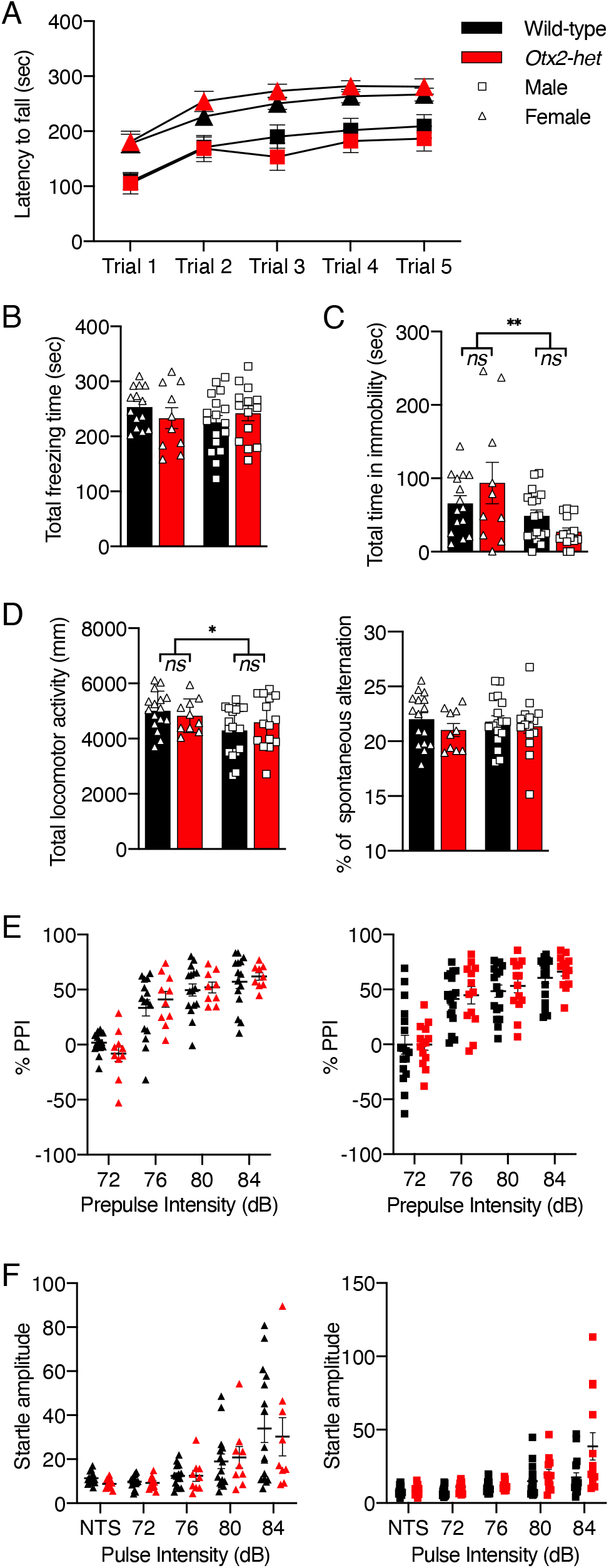
Behavioral analysis of male and female *Otx2-het* mice between P90 and P120. **(A)** Rotarod test with a progressively accelerating rotating rod from 2 to 40 rpm during 5 min trial (WT females n=16; WT males n=18; *Otx2-het* females n=10; *Otx2-het* males n=14). **(B)** Forced swim test during a 6 min period in a small pool (WT females n=16; WT males n=17; *Otx2-het* females n=10; *Otx2-het* males n=14). **(C)** Tail suspension test during a 6 min period (WT females n=16; WT males n=17; *Otx2-het* females n=10; *Otx2-het* males n=15). **(D)** Y-maze test during a 10 min period (WT females n=16; WT males n=17; *Otx2-het* females n=10; *Otx2-het* males n=15). **(E,F)** Prepulse inhibition (PPI) test performed in a startle chamber to measure startle response amplitude (F) and %PPI (E) at various pulse intensities (WT females n=16; WT males n=18; *Otx2-het* females n=10; *Otx2-het* males n=14). All values: mean ± SEM; two-way ANOVA, post hoc Tukey test; *p < 0.05, **p < 0.01.

